# A trans-omics comparison reveals common gene expression strategies in four model organisms and exposes similarities and differences between them

**DOI:** 10.1101/2020.09.04.283143

**Authors:** Jaume Forés-Martos, Anabel Forte, José García-Martínez, José E. Pérez-Ortín

## Abstract

The ultimate goal of gene regulation should focus on the protein level. However, as mRNA is an obligate intermediary, and because the amounts of mRNAs and proteins are controlled by their synthesis and degradation rates, the cellular amount of a given protein can be attained following different strategies. By studying omics datasets for six expression variables (mRNA and protein amounts, plus their synthesis and decay rates), we previously demonstrated the existence of common expression strategies (CES) for functionally-related genes in the yeast *Saccharomyces cerevisiae*. Here we extend that study to two other eukaryotes: the distantly related yeast *Schizosaccharomyces pombe* and cultured human HeLa cells. We also use genomic datasets from the model prokaryote *Escherichia coli* as an external reference. We show that CES are also present in all the studied organisms and the differences in them between organisms can be used to establish their phylogenetic relationships. The phenogram based on 6VP has the expected topology for the phylogeny of these four organisms, but shows interesting branch length differences to DNA sequence-based trees.

The analysis of the correlations among the six variables supports that most gene expression control occurs in actively growing organisms at the transcription rate level, and that translation plays a minor role in it. We propose that all living cells use CES for the genes acting on the same physiological pathways, especially for those belonging to stable macromolecular complexes, but CES have been modeled by evolution to adapt to the specific life circumstances of each organism. The obtained phenograms may reflect both evolutionary constraints in expression strategies, and lifestyle convergences.

## Introduction

The Central Dogma of Molecular Biology states that information runs from DNA to protein [1]. The information flux for protein-coding genes has an obligate intermediary: mRNA (Figure 1). The regulation of the expression for these genes ultimately addresses the control of protein levels in the cell because the final goal is readily protein availability. In fact, protein abundance (PA) seems to correlate much more between different organisms than mRNA abundance (RA; [2]). However, given the central position of mRNA, and because both RA and PA are controlled by synthesis and degradation rates (Figure 1), the desired PA can be obtained following different strategies [3] that balance the contribution of productive and destructive steps, as well as the relative importance of transcriptional and translational regulation [4]. Recent evidence shows that transcription, and not translation, determines PA under steady-state conditions from yeast [5,6] to mammals [7,8]. Moreover, several studies have suggested that changes in mRNA levels in dynamic scenarios strongly determine protein dynamics (discussed in [9]). This topic, however, is open to discussion [4,10–12]. In fact, it has been argued that under severe pleiotropic stress conditions, the contribution of protein-level regulation, translation rate (TLR) and protein stability (PS), is more important. Hence the relative contribution of mRNA-level and protein-level regulation can be context-dependent [4]. It is clear that PA depends on the dynamic balance among these processes, but how this balance is achieved and to what extent all these processes contribute to the regulation of cellular PA are still open questions.

**Figure 1.**
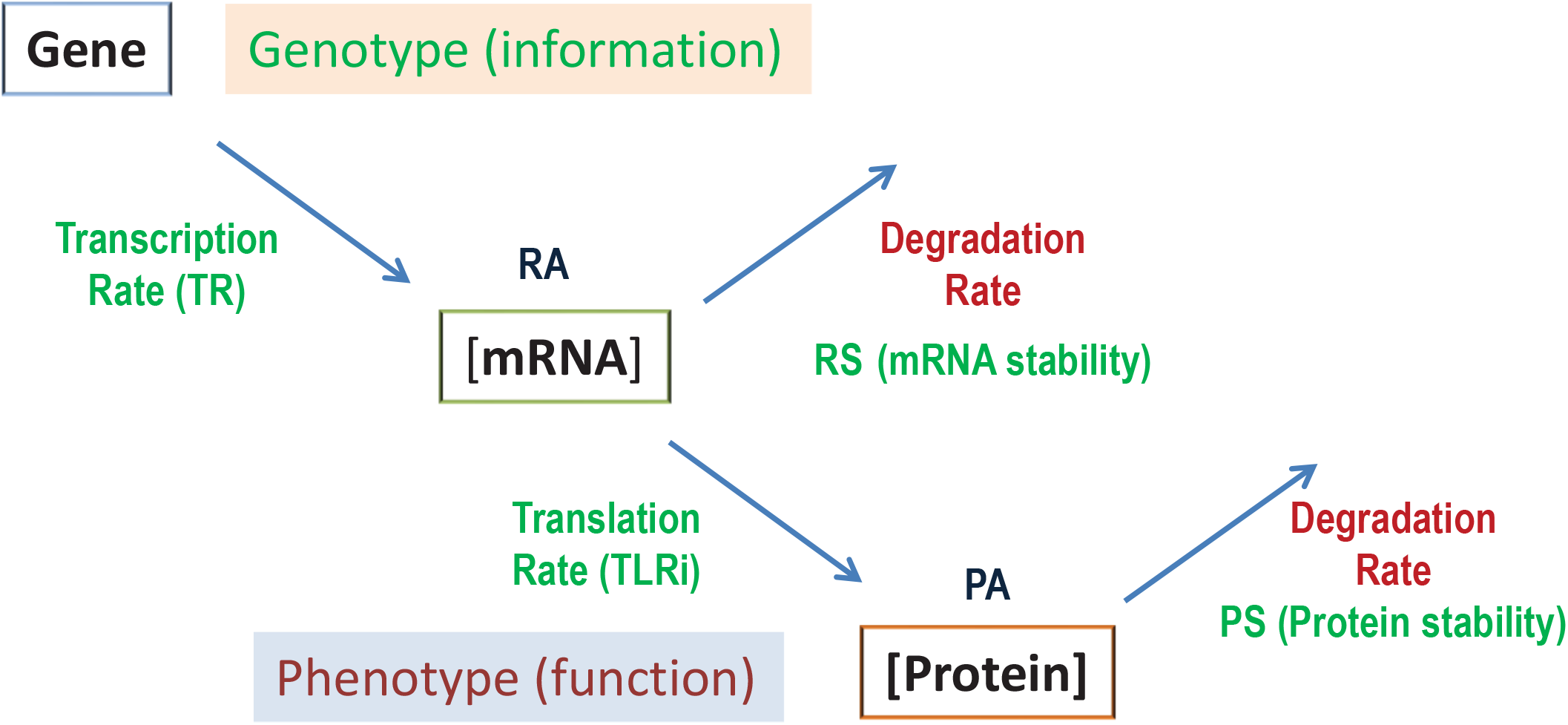
The six variables of the gene expression flux. The genetic information (genes = genotype) is transcribed to mRNAs, whose concentration (RA), under steady-state conditions, depends on the equilibrium between synthesis (TR) and the degradation rates (herein expressed as the reverse parameter, mRNA stability: RS). The phenotype is, however, dependent mainly on the protein concentration (PA) that, in turn, depends on their synthesis (TLRi) and degradation rates (or stabilities, PS). See the main text for additional explanations.

The evolution of extant cells should have taken into account the energy costs of each step. Proteins are thousands of times more abundant than mRNAs and have a larger dynamic range [3,10] that makes their synthesis and regulation much more costly processes. This may be the reason for selecting gene expression mechanisms at the mRNA level [13]. Other variables that influence the selection of specific strategies of gene expression are appropriate speediness and the required level and gradation of the response to potential changes in the environment [14], the optimal biological noise associated with each step [15–17] and the feasibility of post-transcriptional and/or post-translational regulatory mechanisms [16].

In a previous study conducted with the model yeast *Saccharomyces cerevisiae,* we addressed these questions by comparing omics data for the abundances of mRNAs and proteins, and their synthesis and degradation rates (studied herein as their reverse variable: stabilities) [3]. We found that yeast cells use common expression strategies (CES) for the genes belonging to the same physiological pathway. Thus we defined a 6-Variable Profile (6VP) for each functional group of genes to illustrate the particular average expression strategy followed by it. Our results also showed that synthesis rates and molecule amounts tend to have higher correlations between one another than with stabilities, which suggests a more important role for synthesis rates in expression regulation.

In the present study, we check if the results obtained in budding yeast were general to other organisms. To answer this question, we selected three additional model cells for which omics technologies have obtained enough data of the six variables: TR, RS, RA, TLR, PS, PA. We chose another single cell eukaryote, the fission yeast *Schizosaccharomyces pombe,* because of the high quality of the genomic data obtained for it, and also because it is distantly related from budding yeast, but has converged with it in lifestyle [18–21]. As a higher eukaryotes model, we selected HeLa cells because it is the human cell line with the best and most extensive omics data [22]. Finally, we chose the model bacterium *Escherichia coli* as being representative of prokaryotes for similar reasons. Our results show that all kinds of organisms have common expression strategies for functionally-related gene groups and that CES are especially robust for genes coding for subunits of stable macromolecular complexes. CES have similarities and differences between organisms that allow phenetic dendograms (phenograms) based on them to be constructed which, to a great extent, recapitulate the evolutionary tree. There are, however, interesting features which point that functional divergence is not always strictly proportional to sequence divergence. Finally, our results support the notion that most gene regulation takes place at the mRNA synthesis level, whereas translation plays a minor role, but serves to potentiate the effect of transcription.

## Materials & Methods

### Selection and features of the original data

We used data from several publications that have followed, in many cases, different methods or experimental setups. In all cases, data were obtained from the standard reference strains of the four organisms. We sometimes used only one dataset. In other cases in which data from more than one study were available for one variable, the following strategy was followed: first, we compared the datasets by making plots between them. If the Pearson correlation was good (>0.3), we used them all. Second, the data groups corresponding to the same variable were stored in an array and the Bioconductor package was used to impute the unrepresented data by the closest k-neighbors method with the parameters included by default. Then the medians of data groups were matched. Finally, the average was calculated between the values represented for each data group to obtain the values of the final expression, synthesis rate or stability for that variable. The actual data employed for the comparisons are shown in Supplementary Table S1.

For *S. cerevisiae* (S288c background), we updated the datasets previously described used by our previous study [3]. RS was counted with the data from two sources [23,24] obtained by the Dynamic Transcriptome Analysis (DTA) and the RNA Approach to Equilibrium Sequencing (RATE-seq), respectively. Despite not having a large correlation (R = 0.31), they were combined because, although RATEseq technology is the most reliable in methodological terms, the study with DTA included the RA and TR data from the experiment, which improves correspondence (decreases noise) compared to other parameters. In this way, a final list of 5667 genes was obtained. For the RA data, we used the data of [24] and [25], with the technologies based on DNA microarrays and RNA-seq, respectively. They were combined to give a file with 4985 analyzed genes. The PS data were obtained from [26], acquired by Pulse SILAC, and corresponded to 3801 genes. For PA, we used the data from [27], who determined the expression data of 3539 genes by a single cell image analysis of GFP-fusion collection with green fluorescent protein (GFP). The TR data came from averaging the results of [28] obtained by the Genomic Run-On (GRO) method, and those of [23], acquired by the DTA method, which gave 5531 genes. Finally, the employed individual translation rate data (TLRi), collected from the work of [29], which employed ribosome profiling and acquired data for 4623 genes. We used the gene identifiers described in SGD (http://www.yeastgenome.org/).

For the *S. pombe* data, we utilized the following data. For the RS, the data obtained by thiouracil metabolic labeling came from [30] (DNA microarrays), [31] (RNAseq) and [32] (DNA microarrays). Data were combined for a total of 5059 genes. For RA, data were obtained from [32] (DNA microarrays), [33] (RNAseq) and [34] (DNA microarrays). Data were combined for 4800 genes. For TR, data were obtained by thiouracil metabolic labeling and from [32] (DNA microarrays), [31] (RNAseq) and [34] (DNA microarrays) for 5048 genes. The TLRi data were collected from [634] (polysome profiling and DNA microarrays) for 3586 genes. The PA data were obtained by mass spectrometric spectra from [33,35] for 3328 genes. Finally, the PS data were acquired by Pulse SILAC from [26] and corresponded to 2947 genes. We used the gene identifiers described in *pomBase* (https://www.pombase.org/).

For HeLa cells, we employed the following data. For RS, the data were obtained from BRIC-seq for 10817 available genes [36]. With RA, the RNAseq data were available from three sources [36–38], which were combined to give a final list of 18301 genes. The PS data came from analyzing the Chromatography-Tandem MS (LC-MS/MS) of the cells treated with cycloheximide (CHX) with information about 4051 genes [39]. For PA, data came from three sources that correlated well (R>0.3). As what actually matters is the relative expression of each protein compared to the others, although two of them presented expression data as number of protein copies per cell (PCN) [40,41] and the other as iBAQ intensity [37], the last one was converted into PCN with a conversion factor obtained from the plot between them. As a result, a final list with 11269 genes was obtained. Finally, there is no published data on HeLa TR and TLRi data, which were calculated mathematically from the quantity and synthesis data using Equations #1 and #2 (10817 gene data for TR and 3768 for TLRi):

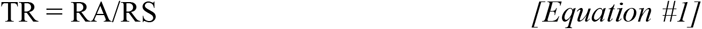

TLR = PA/PS and TLR = RA * TLRi hence:

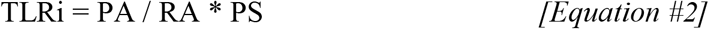

The gene identifiers were converted into the HUGO nomenclature (http://www.genenames.org/) because it is the most widely used one among different sources.

For *E. coli,* we used the following data obtained from the K12 strain exponentially grown at 37°C in M9 or LB media. For RS, we used the data from [42,43] obtained by transcription shutoff with rifampicin and DNA microarrays. They were combined obtaining a final list of 2947 genes. For RA, we utilized the datasets from [42,43] (DNA microarrays), [44] (RNAseq) and [71] (DNA microarrays). They were combined to give a final list of 4284 genes. For TLRi, we used the data from [44] (ribosome profiling) that corresponded to 2387 genes. For PA, we employed the data from [43] (SILAC), [45] and [46] (mass spectrometric spectra). They were combined to give a final list of 2045 genes. The TR and PS data were obtained mathematically from other data using Equations 1 & 2 to give a list of 2947 and 1928 genes, respectively. We used the gene identifiers described in the EcoCyc Database [47].

### Correlation analyses

To test the global correlation among all the pairwise combinations of the six variables (obtained as explained in the previous section), Pearson’s correlation coefficients (R) and their associated p-values were calculated using the data from the genes for which complete information was available.

Then by using this same information, a Bayesian Model Averaging (BMA) procedure [48] was performed using BayesVarSel R package [49]. This approach allowed us to make a robust estimation of the coefficients of each variable in a multiple linear regression to explain PA. It worth mentioning that a BMA estimation of a coefficient takes into account the potential correlations among the variables included in the linear regression by weighting each possible combination of them according to its corresponding posterior probability. Note that, in order to make all the coefficients comparable, data were transformed using Z-scores. Accordingly, Figure 2B presents the posterior mean and 95% credible interval for each coefficient and organism. Certain variables were not included in the analysis for some organisms given their direct mathematical calculation from the response variable (PA), which was the case for TLRi in HeLa and PS in *E. coli*.

**Figure 2.**
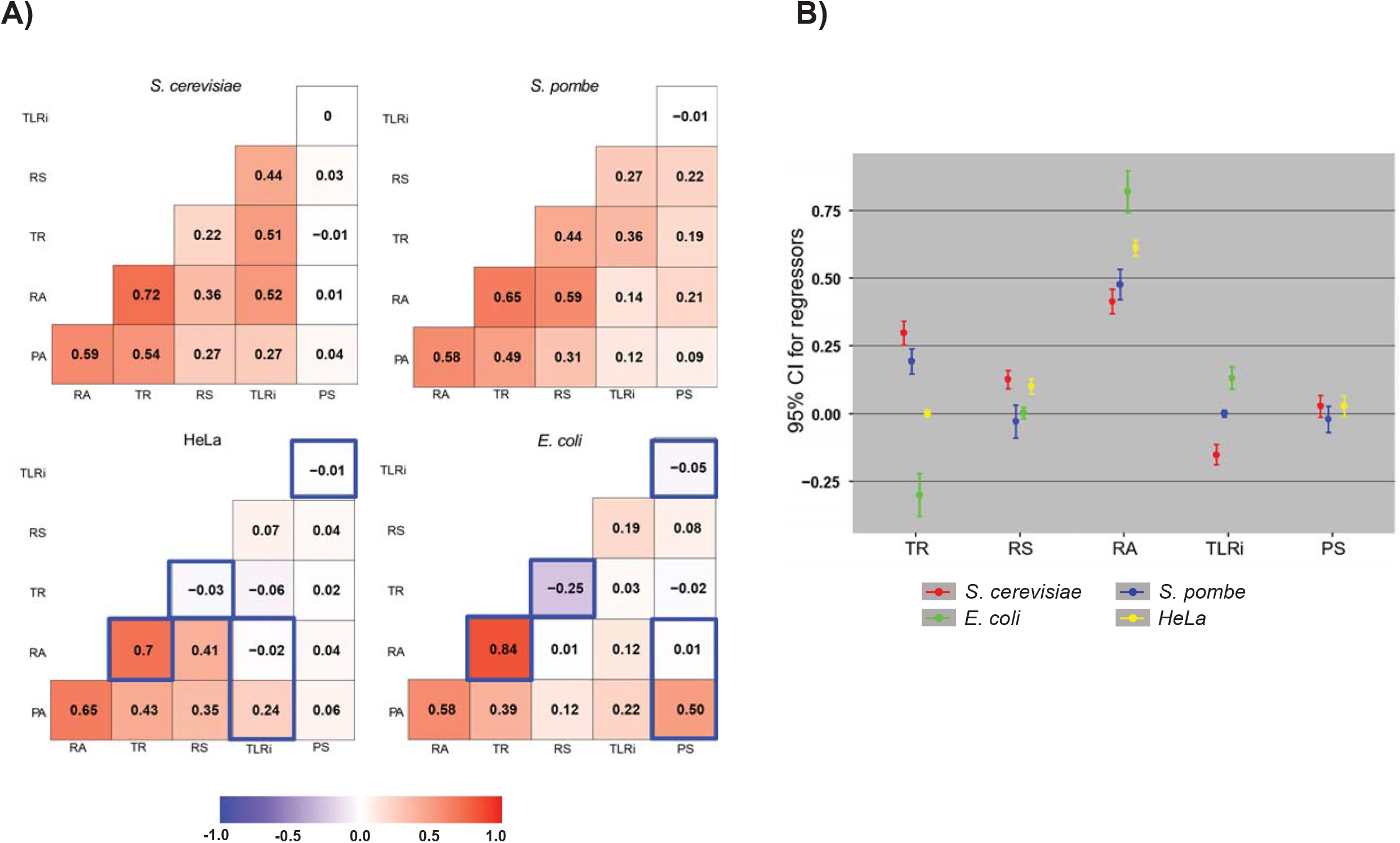
Correlations between the six variables of the gene expression in four model organisms. A) The pairwise Pearson’s correlations among the six variables in each organism. Exact values are indicated in frames. The background color scale indicates positive (red) or negative (blue) correlations. Some correlations are less reliable because they are between the variables calculated from some others in HeLa and *E. coli* (see the main text) and are marked with blue boxes. B) Estimation of the coefficients of a multiple regression model per organism using Bayesian Model Averaging (posterior mean and 95% credible interval). These coefficients show the contribution of each variable to the final protein amount (PA) using a z-score scale to make them comparable. The TLRi for HeLa and the PS for *E. coli* are not shown because they were mathematically calculated from PA values (see M&M).

### Six variable profiles (6VP)

We analyzed the behavior of the genes from all four organisms using rank data values (0 was the lowest value, and 1 was the highest (Fig. 3A) to avoid the wide dispersion in the unit ranges seen when comparing the different datasets for the six variables obtained with very distinct experimental techniques. In this way, although some information was lost, the results were much more robust. Matrices were obtained with 3613, 4139, 3350 and 1643 genes represented for HeLa, *S. cerevisiae*, *S. pombe* and *E. coli*, respectively (Supplementary Table S2).

**Figure 3.**
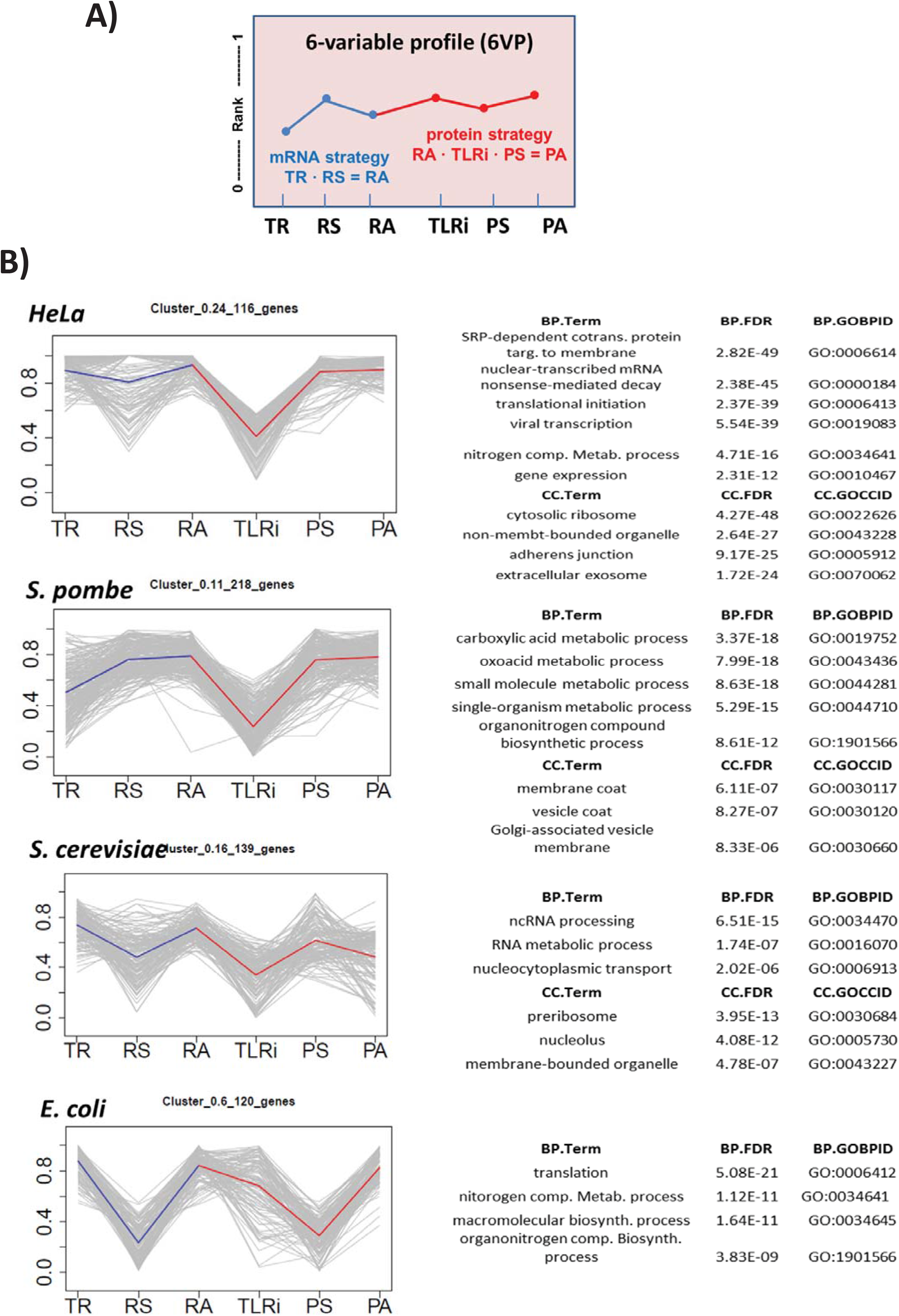
The 6-variable profile (6VP) allows to cluster all genes according to their expression strategy. A) Genes were ranked from lowest (0) to highest (1) in each variable (TR, RS, RA, TLRi, PS, PA). The shape of the line linking the six points is characteristic of an expression strategy: 6VP. The strategy to gain a defined RA corresponds to the first part of the line (in blue): in the steady state the mRNA level depends on both its transcription rate and stability. For the protein strategy (in red), the final level depends on both the total translation rate (TLRi x RA) and protein stability. B) Many clusters selected from the four studied organisms show defined profiles and statistically enriched GO terms. The individual gene profiles are shown in gray and the average profile of the cluster on the colored line. Some of the GOs with the highest p-value are shown on the right for the Biological Process (BP) and Cellular Component (CC) ontologies. Other examples are found in Supplementary Fig. 1 and the whole lists are given as Appendices.

The behavior of the genes belonging to the functionally related eukaryotic gene groups was analyzed. We selected some GO terms with enough genes in all three eukaryotes (Figure 4A) or groups of functionally related genes (Figure 4B) that were obtained from a previous selection, described in [3]. The definition of groups was based on *S. cerevisiae* genes. The orthologous genes from the other eukaryotes were obtained from the YeastMine database (http://yeastmine.yeastgenome.org/) for HeLa and from *pomBase* (https://www.pombase.org/) for *S. pombe.*

**Figure 4.**
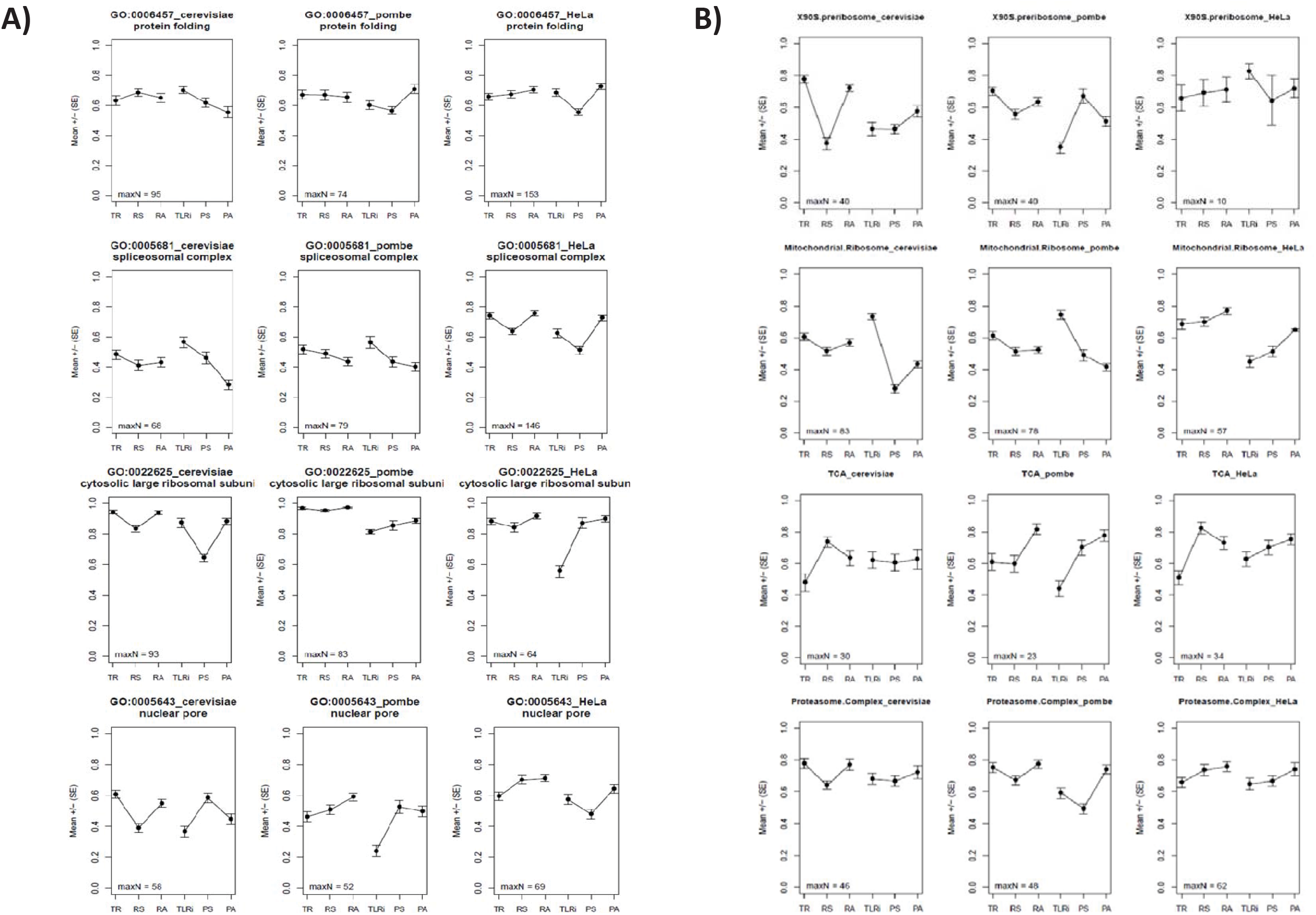
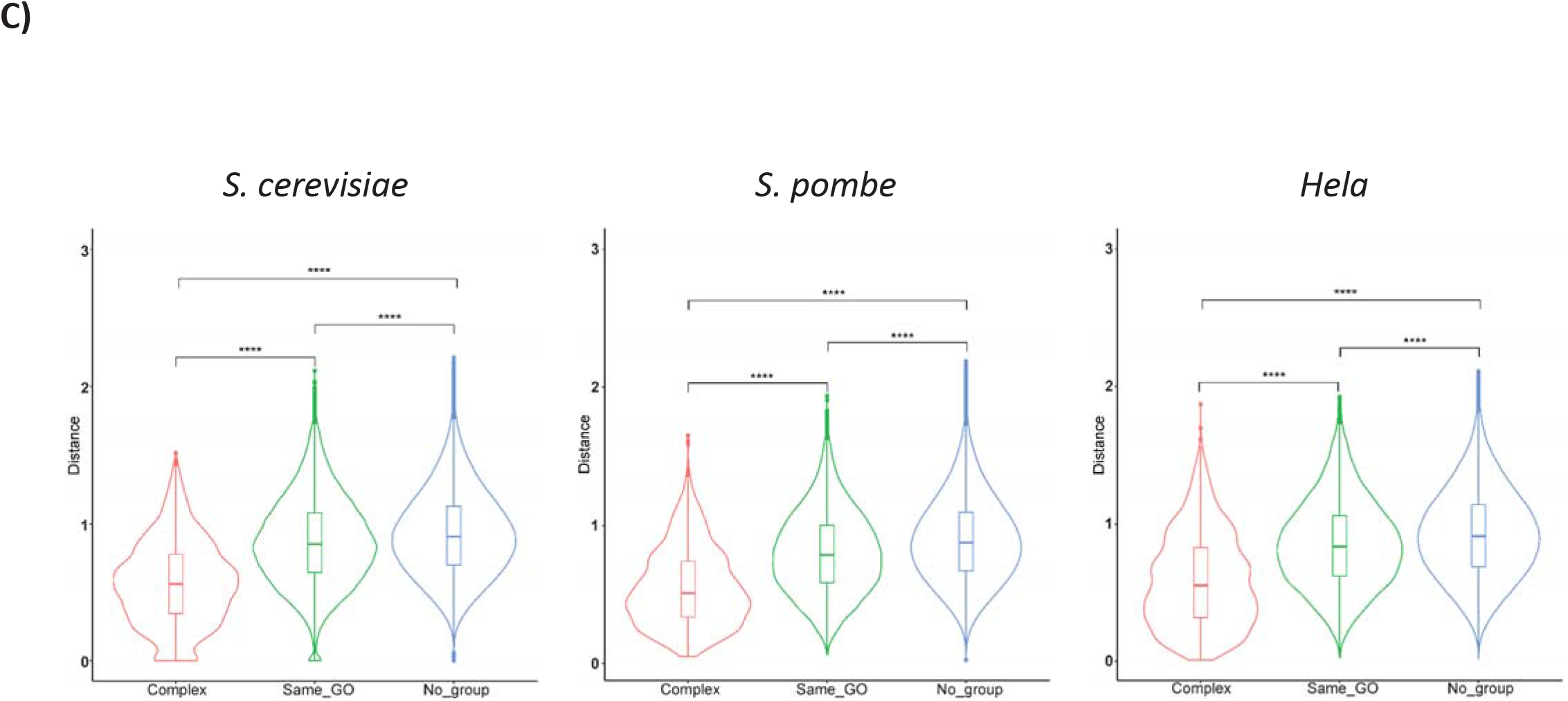
6VP for the selected GOs and manually-curated groups in *S. cerevisiae*, *S. pombe* and HeLa. A) We selected four GO terms corresponding to well-known groups of functionally related genes/proteins. The ID of the GO and its name are shown for *S. cerevisiae* (left), *S. pombe* (center) and HeLa (right). The maximum number of genes used (*maxN*) to determine the average value and standard error is shown. In some variables, this number may be lower due to lack of information for some particular genes in some datasets. In this figure, unlike Figure 3, we independently represent mRNA and protein parts without a connecting line to better display the differences between them. B) A similar analysis to that in A), but done with the manually-curated categories that were used in a previous study in *S. cerevisiae* [3]. C) Violin plots showing the mean of proximity between the genes coding for the proteins acting in macromolecular complexes (Complex, red) belonging to the same GO, but do not form complexes (Same_GO, green) and other gene pairs that do not belong to the previous groups (No_group, blue). A t-test was applied for the statistical significance of the difference. *** means p-value < 2.2e-16 (lowest numerical value displayed in R). An ANOVA also detected a significant difference (p-value < 2.2e-16) among the three gene groups in all the organisms.

We calculated the average rank value and represented these values for the six variables in this order, TR, RS, RA, TLRi, PS and PA, to yield a 6VP for each studied group. We also calculated the standard error (SE) associated with each average and represented it in the profile as error bars. A control test was done by averaging values of 1000 random samplings with the same sample size than the analyzed functional group. In all cases they appear as a flat line at the 0.5 score. They have been omitted in the graphs for clarity.

### Cluster analyses

In order to identify the groups of genes with similar expression profiles, gene clustering was done according to their 6VP (Fig. 3). In this way, we settled characteristic expression profiles with the data of the six variables for the genes with at least four represented variables (of which at least two had to be of mRNA and the other two of protein).

For the clustering analysis, the *sota()* function of the R *clValid* package [50] was followed. This function performs clustering with the *Self Organizing Tree Algorithm* (SOTA) [51] using the linear correlation coefficient among the six variable vectors as the distance between genes, following a splitter scheme that allows the algorithm to be stopped at any point to gain the desired number of clusters. This algorithm does not allow any variables with missing data, so the Bioconductor impute package was used to impute unrepresented data by the closest k-neighbors method. The algorithm was settled in order to avoid clusters with less than five genes.

Trees were allowed to grow until 10, 15, 20, or 30 clusters were produced. Then clusters were manually selected from any clustering level by considering the p-value of the enrichment for GO terms by looking for the best ones, and in such a way that clusters do not overlap.

### Gene Ontology category searches

To test potential enrichment in GO terms in the different clusters obtained by SOTA, and as previously explained, we used the *GOstats* R package. For this purpose, a test based on hypergeometric distribution was run for the terms or divisions of ontologies Cell Component (CC) and Biological Process (BP). Only GO terms were considered significant when applying the *Multitest Correction False Discovery Rate* (FDR) method [52] and they had an adjusted p-value of <0.001. These terms were filtered by a semantic comparison process with the help of the *GOSemSim* package. Using this package, a function was designed to select the GO group with the lowest adjusted p-value of all those with a semantic similarity greater than 70%.

### Comparison of the proximity in 6VP among the genes belonging to macromolecular complexes and those belonging to GO categories non forming macromolecular complexes

We selected 18 stable protein complexes in *S. cerevisiae* that have more than five and less than 150 genes from the MIPS database described in the previous study [3]. We also selected 15 GO categories that include less than 350 genes and are known to not include complexes. Supplementary Table S3 indicates the selected GO terms and the genes included in those for which complete data were available. The *S. cerevisiae* distance matrix included information about 2592 gene pairs that participate in the same protein complexes (Complex), 33558 gene pairs placed in the same selected GO categories with no protein complexes (Same_GO) and 3537651 gene pairs which were not included in either group (No_group). The mean pairwise distance for the genes included in protein complexes was 0.597, whereas these distances were respectively 0.876 and 0.924 for the gene pairs included in the same GO category and the gene pairs not included in either group. The ANOVA showed significant differences between groups and the *post hoc* t-test determined that all the pairwise tests were significant. With *S. pombe,* after the orthologous conversion the distance matrix included information on 2396 gene pairs that participate in the same protein complexes, 15279 gene pairs placed in the same selected GO categories and 1339453 gene pairs not included in either group. The mean pairwise distances for the genes included in protein complexes was 0.554, whereas they were respectively 0.803 and 0.895 for the gene pairs included in the same GO category and the gene pairs not included in either group. The numbers for Hela were as follows: 2660 for the gene pairs in complexes, 8512 for the gene pair distances in the same GO categories, and 793374 gene pairs for the No Group class. The mean pairwise distances for the genes included in protein complexes was 0.591, whereas they were 0.855 and 0.923, respectively, for the gene pairs included in the same GO category and the gene pairs not included in either group.

### Phenogram tree construction

We performed both neighbor-joining and hierarchical clustering using the information deriving from our GO term level 6VP as input. The analysis was performed as follows: first, we removed excessively broad GO terms and the GO terms containing very few genes. Thus in order to keep a functional category for the downstream analysis, it had to contain a number of genes between 5 and 275 in all four species. We only included those functional categories for which we had data about all six variables.

We employed only Biological Process (BP) and Cellular Component (CC) ontologies for searches. In order to remove very close related GO terms, which would probably present large gene overlaps, we followed a procedure inspired by REVIGO [44]. First, we used the *GOSemSim* package [53], a package designed for the semantic comparisons of Gene Ontology (GO) annotations. All the analyses were carried out by taking the human GO database as a reference. From our list dataset of the GO terms, we selected those found in the human GO database. Then for the list of retrieved GO terms, we computed a matrix of pairwise similarity values by the *Rel* method. In short, the *Rel* method combines Resnik’s and Lin’s methods [54,55] to compute semantic similarity (*Sim_REL_*) between any given pair of GO terms. *Sim_REL_* values range from 0 to 1, and the higher the value is, the greater the similarity between GO terms. The pairwise matrix of the *Sim_REL_* values were transformed into a distance matrix (*Sim_REL_-dist*) by computing 1-*Sim_REL_*. For each functional category, a parameter called uniqueness was computed as 1 minus the average of the *Sim_REL_* values of each GO term to all the other terms. This parameter indicates how different a specific GO term is compared to all the others.

The distance matrix, *Sim_REL_-dist*, was then employed to perform hierarchical clustering by the average (UPGMA) method. The mean silhouette information was extracted for any possible divisions from 1 to the number of the included functional categories −1. The number of clusters yielding the highest average silhouette value was selected. Then for each cluster of the GO terms, the term with the highest uniqueness value was selected as the most representative element of each cluster. For those clusters with only one element, this one element was selected. The set of representative elements of each cluster included the GO terms used in the downstream analysis. Lists of the GO terms employed in each tree are shown in Supplementary Table S4.

Using the whole set of GO terms yielded by the previous procedure, we carried out a clustering analysis by two different methods: hierarchical average (UPGMA) and Neighbor-joining employing the *NJ* and *hclust* methods implemented in *phangorn* [56] and *stats* packages (Fig. 5A). We also visually inspected the GO terms and created groups of GO terms linked with the following cellular processes: Cytoplasmic translation, Mitochondrion, Transcription and Replication to make the clustering analysis of selected terms shown in Figure 5B.

**Figure 5.**
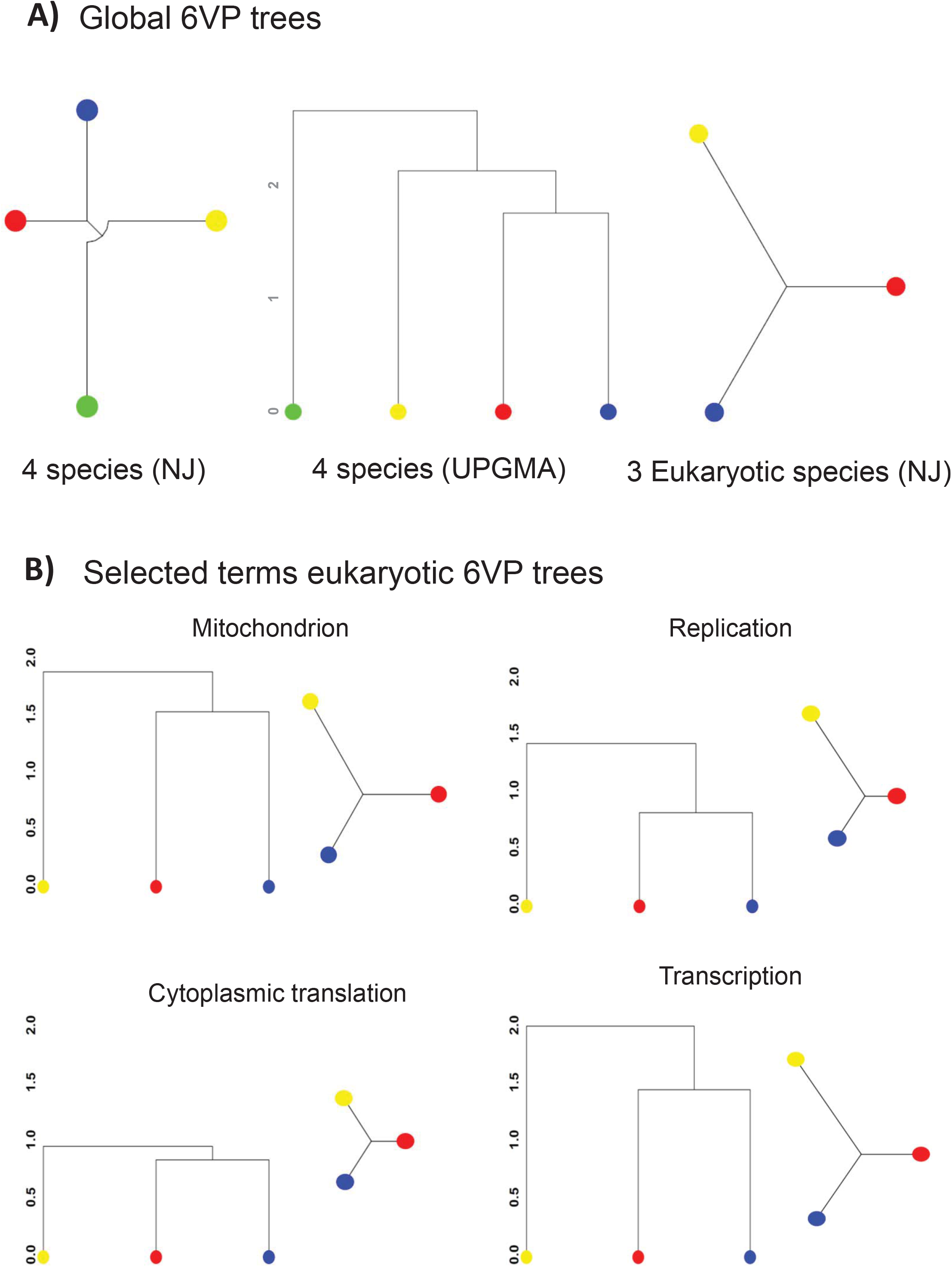
Phenograms of the four organisms based on their 6VPs. A) Global Neighbor-joining (NJ) and UPGMA trees based on the 6VP derived from 53 GO biological process (BP) and 2 GO cellular component (CC) terms for *S. cerevisiae* (red dots), *S. pombe* (blue dots), *E. coli* (green dots), and *H. sapiens* (HeLa cell line, yellow dots) (left and center), or only the NJ tree for the three eukaryotes with 353 GO BP, plus 84 GO CC terms (right). B) NJ and UPGMA trees based on the available GO terms linked (manually curated) with cytoplasmic translation, transcription, mitochondrion and replication. All the NJ and UPGMA unrooted trees are, respectively, on the same scale, but NJ ad UPGMA are represented with a different kind of branching to show complementary information. The lists of the GO terms used in each tree are provided in Supplementary Table S4.

## Results

### Datasets for the variables selected in this study

In this work, we studied the six variables that control gene expression (see Figure 1)transcription-rate (TR), mRNA amount (RA), mRNA stability (RS), translation rate per mRNA (TLRi), protein amount (PA) and protein stability (PS)- using omics datasets under a standard growth condition for each studied organism. When the published work studied different growth conditions, we selected that with the highest growth rate. Although the right parameter to be used is concentration rather than amount, we can assume that the cell volume for each organism is constant for all their datasets obtained under the same culture conditions and, thus, variations in the number of molecules and their concentrations are equivalents As the large differences within the ranges of actual values among the six variables we used ranks and Z-scores instead of absolute values to make the results more robust.

In a previous study into *S. cerevisiae* [3], we used the datasets available for the omics data at that time. We have updated some of the datasets from *S. cerevisiae* by taking special care with the mRNA half-lives dataset that produces very different results from the previous one (see below). For *S. pombe*, HeLa and *E. coli* we selected from the available datasets. When two datasets or more were available, we took the average of the well-correlated ones to be the final value (see M & M for a detailed protocol). For human cells, we selected the HeLa datasets because this cultured cell line covers more information about omics data. Although other human cells lines in culture may evidently have different data for specific genes, we assume that the global behavior in HeLa cells is the best available possibility and is a representative of human cells in culture.

The actual synthesis rates of mRNAs and proteins, TR and TLR are, in fact, the product of individual rates, namely TRi and TLRi, multiplied by the number of genes or mRNA copies, respectively. For mRNA synthesis rates, in practice TR and TRi are equivalents for single cell organisms because most genes have one (in haploids) or two (diploids) copies. Hence we used the acronym TR throughout this paper. Although HeLa cells are considered mostly diploid, they have a high aneuploidy level and numerous large chromosome structural variants [57]. Therefore, the TR in HeLa is not equivalent to TRi, but is the right value to be used because is the actual determinant of mRNA levels. For protein synthesis, TLR and TLRi are, however, essentially different. Given its dependence on RA (see M & M), TLR is mathematically linked with it. Yet TLRi reflects an intrinsic property of mRNA and has been calculated experimentally, usually as ribosome density, by ribosome profiling [29]. For this reason, we employed TLRi values for our analyses.

Finally, it should be noted that the quality of the datasets is not the same for the four organisms. These kinds of omics studies have been conducted more frequently in yeasts. The gene coverage in both yeasts for most variables is much better than for HeLa and *E. coli*. The existence of mRNA and protein isoforms also adds a complication to the interpretation of the study. The number of this isoforms is higher in HeLa. In this study we have grouped all isoforms present in datasets under a single gene name. The number of datasets for both yeast species is much bigger and we could select several and make an average dataset. Moreover, the yeast datasets for all variables consist in direct experimental data. For HeLa and *E. coli*, however, some variable datasets were not available and we had to estimate them mathematically from others (see M & M). Therefore, we consider that our conclusions are more robust when taken from budding and fission yeast variables.

### Comparisons and correlations between variables

It is believed that protein amounts depend mainly on mRNA amounts under steady-state conditions [4,8], although the correlation value is still a matter of discussion [6,58,59]. Thus a positive correlation between RA and PA datasets is expected. In fact, this has been previously observed for the three eukaryotes herein studied [3,33,37,60]. Other correlations have been much less studied [3,6,9], but may illustrate the strategy adopted by cells to determine a given PA.

In this study we obtained pairwise correlations (Pearson) among the six variables considered for each organism (Figure 2). It is worth noting that the actual correlations between variables may be underestimates of the true correlations due to measurement errors in the employed datasets. For all the organisms, we found positive and statistically significant correlations (red backgrounds in Figure 2A) between RS and TR with RA, which means that both synthesis and degradation rates (i.e. mRNA stabilities) control mRNA levels. The higher TR/RA correlations indicate a stronger influence of TR on mRNA levels. RAs also positively correlate with their stabilities, and take lower values, in the three eukaryotes, but not in *E. coli*. This argues that stabilities are not as important as synthesis rates for determining mRNA levels. In our previous work [3], we found a negative correlation between RS and RA in *S. cerevisiae*. This time we used new more reliable RS datasets, and this result changed. We think that, together with other authors [24,32,61–66], the RS datasets obtained by transcription shutoff methods are affected by marked biases that invert the observed correlation [61]. In *E. coli,* we found no correlation or a negative correlation between RS and RA or TR, probably due to the RS dataset having been obtained by transcription shutoff [42].

The large correlations between RA and PA, and the lesser correlations between PA and TLRi, argue that, as recently suggested [5–8], mRNA levels are much more important for determining protein levels than specific translation rates, although TLRi can be very different between mRNAs and explain part of the actual PA level [6]. However, it should be taken into account that, for HeLa, the TLRi values were derived as a mathematical calculation involving PA, RA, and PS (see M & M) and, therefore, the corresponding correlations (marked with a blue box in Fig, 2A) may be influenced by this fact. In any case, the message of the low positive, but statistically significant, TLRi-PA correlation in the other three organisms supports the idea that abundant mRNAs tend to be better translatable. This is probably because they are also enriched in optimal codons [67], which has been called potentiation or amplification exponent [5,8]. The positive correlations of TLRi seen in the yeasts with RA, TR and RS also support the idea that a potentiation effect occurs during translation. Following the same argument, we can conclude from the TLRi-RS positive correlation that a direct proportionality appears in free living cells (at least in eukaryotes) between the stability of an mRNA and its capacity to be translated, which confirms the results from J. Coller’s [68] and other laboratories [61] by showing that in *S. cerevisiae* mRNA enriched in optimal codons (better translatable) are more stable than those depleted in such codons. Similar results have been obtained for *S. pombe* [69] in *E. coli*, zebrafish and mammalian cells (reviewed in [67]). Our analysis cannot confirm this for *E. coli* and, especially HeLa, which can be due to a genuine lack of correlation or to an artefactual bias due to the direct mathematical calculation of some variables as mentioned above).

The existence of much better correlations between TLRi and PA than between PS and PA argues that for protein amount, and even more strongly than for the mRNA amount, the total synthesis rate (TLR, e.g. multiplying RA by TLRi) is the main determinant. In fact it would seem that protein degradation is not used by any organism as a way to determine the most steady-state protein levels. This is not surprising for fast living single cells in which protein half-lives are usually much longer than generation times, which implies that dilution by cell division is a much more important factor for protein disappearance than protein degradation [13]. Moreover, for all organisms, including HeLa cells where the generation time is longer, the high energy cost of regulating abundant proteins by degradation does not seem to be a suitable strategy [13].

In order to test the contribution of each variable to the final protein amount (PA), we performed a multiple regression analysis based on a Bayesian Model Averaging approach [49] to evaluate what proportion of the final PA is due to each variable in the four studied organisms. The result is shown in Figure 2B. It is clear that for all four organisms, RA is the most important contributor to PA. In the two yeasts, TR also makes a significant contribution. In *E. coli* and HeLa, as previously explained, TR was mathematically calculated and is less reliable. The other three parameters, RS, TLRi and PS, contribute much less to PA.

These results support the conclusion that the main determinant of the protein level in a cell is the corresponding mRNA level and that RA, in turn, depends mostly on the transcription rate of the gene.

### Clustering of genes according to the six variables of gene expression into four different organisms

Our previous results obtained with *S. cerevisiae* demonstrated that functionally related genes tend to be grouped according to their gene expression variables [3]. In this work, we repeated the clustering for that yeast with new datasets, and for the other studied three organisms using the previously described omics datasets. Here, however, we used ranked values (see Supplementary Table S2) because the ranges for the six variables were quite different. We employed the six values for clustering in this order: TR, RS, RA, TLRi, PS and PA (6VP; see Figure 3A). We performed a clustering analysis of the genes for which the data on at least four of the six variables were available (Supplementary Table S2). In this way, we analyzed 4139 genes for *S. cerevisiae*, 3350 genes for *S. pombe*, 1653 genes for *E. coli* and 3613 genes for HeLa cells.

Thus we obtained a 6VP for each gene. This allowed us to compare all the genes for common profiles by standard clustering methods. By this procedure we found that clusters had genes with similar profiles that were statistically enriched in the Gene Ontology (GO) categories (terms) in all four organisms (Figure 3B, Supplementary Fig. 1 and Appendices). This result extends our previous results obtained with *S. cerevisiae* [3], and demonstrates that common expression strategies (CES) for the genes with a related biological function are a common feature of living beings. It should be noted that the quality of the datasets for the four organisms (see above) can influence clustering quality. We consider that the 6VP is more robust for the two yeasts, and in HeLa at a lower level. For *E. coli,* the poorer quality of the datasets and the very poor quality of the information in the GO annotation for this organism [70] precluded the finding of strongly enriched clusters.

### Detailed analysis of the selected functional groups in eukaryotes

If all the analyzed organisms have CES for, at least, part of their gene groups, we may wonder if the particular CES followed by a given gene group is similar or different in distinct organisms. Given the poor quality of the *E. coli* annotations, we compared only the three eukaryotes. Figure 4 depicts some examples of these comparisons. We studied either selected GOs (Fig. 4A) or some of the manually-curated groups (Fig. 4B) analyzed in a previous work in *S. cerevisiae* [3] to look for the orthologous genes in *S. pombe* and HeLa.

It can be seen that the average profiles for the groups generally show a similar ranking for all the variables in the three organisms. For instance, protein folding (GO:0006457) shows that all the six variables rank 0.6-0.8 in all three organisms, whereas cytosolic ribosome (GO:0022625) ranks higher than 0.8 for most variables and the nuclear pore (GO:0005643) ranks mostly between 0.4-0.6. This demonstrates that the particular levels of mRNAs and proteins for a given group of genes tend to be similar in all eukaryotes and, to a lesser extent, that the strategies followed to this end are also similar. It is interesting to note that some others differ. For instance, the spliceosomal complex (GO:0005681) ranks higher in HeLa than in the two yeasts. This result is logical given the much more marked importance of splicing for human genes [71].

Regarding the shape of profiles, we can found similarities and differences. V-shaped profiles (meaning lower stabilities than synthesis rates) are common in stable macromolecular complexes in *S. cerevisiae,* especially for the mRNA part (see Fig. 3A-B). This has been previously noted [3]. This feature has also been noted for human THP-1 and C2C12 cells [472], but is not so common in *S. pombe* and HeLa cells where only some stable complexes behave as such. For instance, cytosolic and mitochondrial ribosomes have V-shaped profiles in budding yeast, but are not so marked in fission yeast and HeLa for the protein part. The spliceosomal complex is more clearly V-shaped in HeLa than in both yeasts but proteasome is, conversely, V-shaped in both yeasts, but not in HeLa. To conclude, we can state that the existence of cases with similar and different strategies in the three model eukaryotes make 6VP suitable for comparing the expression strategies for the whole gene sets between different organisms.

Both this analysis and the previous one in *S. cerevisiae* [3] suggest that the genes coding for proteins that are subunits of stoichiometric stable complexes tend to have better defined 6VP than other functionally-related gene groups. To test this hypothesis, we performed a comparative analysis of the profile distances between the gene pairs belonging to protein macromolecular complexes and the genes belonging to the GO categories not including macromolecular complexes (Figure 4C). It is clear in all three eukaryotes that 6VP distances are much lower between the genes belonging to complexes, although the genes belonging to the same GO category that does not form complexes are still closer than random gene pairs.

### Trans-organism clusters comparison: 6VP phenograms

As we previously observed cases in which 6VP for GO terms were similar between some organisms, but different in others, we wondered if the whole similarity of the 6VPs among the four studied organisms could be used to make a phenetic tree based on the similarities of the expression strategies for the same functional groups among the four organisms. In order to do this, we selected the GO terms with a number of genes between 5-275 to avoid excessively small groups, which can bias clustering, and the excessively broad ones containing not functionally related genes. In this set of GO terms (from Biological Process, BP, and Cellular Component, CC, ontologies), we reduced the redundancy of similar GO terms by applying a procedure inspired in REVIGO pipeline [73] (see M & M). The goal of the redundancy reduction step was to avoid the information about genes present in highly redundant GO terms to excessively influence the following cluster and tree construction.

As we can see in Fig. 5A, the topology of the global Neighbor Joining (NJ) and UPGMA trees is identical to the known topology of the DNA sequence-based tree [19,74]. Given the poor quality of the GO annotation in *E. coli,* the global tree can only use 53 common GO terms for the four organisms (51 BP + 2 CC), and we consider that the branching of this prokaryote is less robust. However, *E. coli* can be considered an outgroup for the eukaryote tree. We repeated the tree using only the three eukaryotes, which extended the set of common GO terms to 437 (353 BP + 84 CC). The topology of this NJ tree is identical and the relative branch lengths are similar to the previous one. The lengths of the branches between the two yeasts and HeLa indicate that *S. cerevisiae* comes slightly closer to human cells in gene expression strategies than *S. pombe*.

We wondered if the topology of the tree and the lengths of branches were the same for the different functional categories of genes. We repeated clustering and tree reconstruction by separately using groups of the GO terms belonging to broad eukaryotic cellular functions, such as those related to macromolecule synthesis processes (Replication, Transcription, Cytoplasmic Translation) or to the Mitochondrion, because this is a large set of genes known to be coordinately regulated [3,75]. In Figure 5B illustrates that the UPGMA trees repeat the topology of the global one. Distances in NJ trees are always shorter between budding yeast and human cells, except for the Mitochondrion group, where the fission yeast comes closer to HeLa. Thus we can conclude that, in terms of cell functions, the two yeasts are much more similar to one another than to human cells, and budding yeast comes slightly closer to human cells than the fission yeast. Finally, the lengths of the branches in both the UPGMA and NJ trees differ for each broad group, which suggests that distinct cell functions and components are more conserved (Cytoplasmic translation) than others (i.e. Mitochondrion, Transcription).

## Discussion

A capital question in Biology is how genetic information is converted into function: how the variable copy number of a given protein is obtained from the constant copy number of its gene. Cells have multiple steps in which the gene expression flux can be regulated. In a series of relevant papers, M. Biggin’s group [8,9] and others [5,6] have shown that protein levels under steady-state conditions are explained mostly by mRNA levels in both yeasts and mammals, and that these levels depend mainly on synthesis rates [9]. In a previous study in the model organism *S. cerevisiae,* we developed a pipeline to compare the respective influences of mRNA and protein synthesis and degradation rates to the final level of most of the proteins of this yeast. We found that the genes belonging to functionally-related groups followed a similar gene expression strategy, which can be defined by a ‘six variable profile (6VP)’ followed by the genes included in it [3]. In the present study, we extend this pipeline to three other model cells to check the extensibility of our previous conclusions.

Our results support the idea that the main pathway used for gene expression is based on synthesis rates for all organisms. The transcription rate is the main determinant in all four organisms of the mRNA level which, in turn, is the main determinant of the protein level (Figure 6). Obviously, this result does not exclude the existence of cases in which other variables also have an effect, or are even the main determinants of protein levels. By using the tRNA modifications enzyme mutants in *S. cerevisiae,* Chou et al, [76] have demonstrated that the number of translationally regulated genes is quite small (57 cases). In another recent study, the careful analysis of RA and TLRi correlations with PA in the same yeast found fewer than 200 proteins, apart from the general tendency of strict correlation between RA and PA [77]. All these results argue that translational regulation is statistically uncommon and is, therefore globally, a minor participant in eukaryotic (and prokaryotic?) gene expression regulation. However, it is worth noting that, in the four organisms herein studied, the data were obtained from actively growing cells. Nonetheless, it is rather possible that the respective importance of the synthesis rates and stabilities vary under no active or very slow growth conditions.

**Figure 6.**
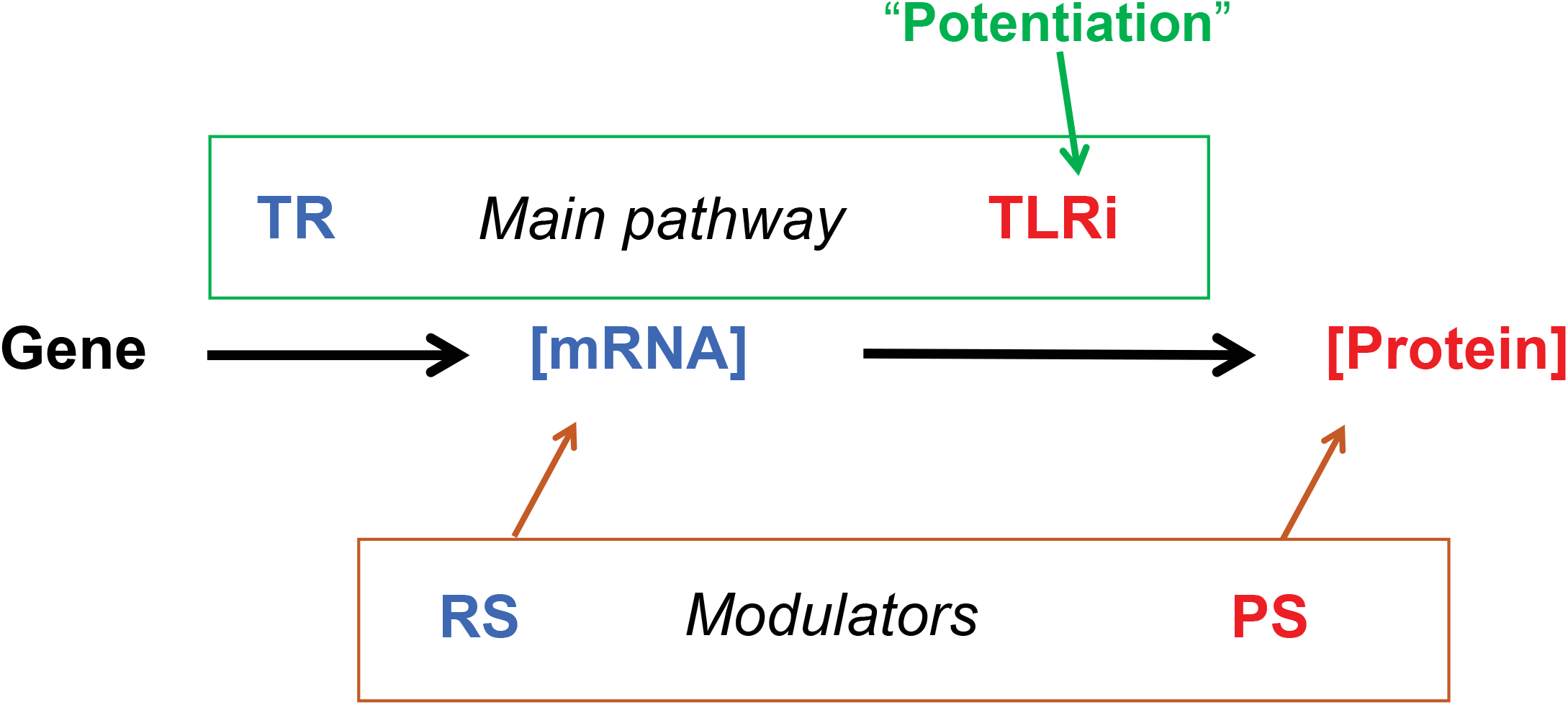
General model for gene expression flux control. In gene expression flux, the transcription rate (TR) is the main determinant of mRNA amount (RA) that, in turn, is the main determinant of protein amount (PA). Therefore, the main pathway for gene expression flux is based in synthesis rates. The individual translation rate (TLRi) of each mRNA potentiates the effect of RA on PA [5,8] because, probably, it depends on the enrichment of optimal codons of mRNAs, which is biased toward abundant mRNAs [67,68]. The control at the stability level of mRNA and proteins has a minor influence globally, but can modulate specific genes or specific situations.

Another interesting topic is the positive correlation of the individual translation rate of each mRNA molecule (TLRi) and the protein level (PA). If TLRi were the same for all the mRNAs, the correlation between RA and PA would still be observed. Therefore, the additional positive correlation of TLRi with RA and PA in all four organisms means that more abundant mRNAs tend to be specifically better translated. This can be explained using the known enrichment of abundant mRNAs in optimal codons that make them more stable and translatable [67–68]. Another possibility would be that transcription imprints mRNAs at a level that depends on the actual TR, in which case mRNAs could be more or less translatable (TLRi) according to their TR. This has been shown to occur in mammalian cells, where methylation of adenines in position 6 (me^6^A) is greater for less transcribed mRNAs and this feature is detrimental for their translation [78]. This cannot be a reason for the potentiation effect on microorganisms because in those organisms, me^6^A is not present in *S. pombe* and *E.coli* and occurs only during meiosis in *S. cerevisiae* [79]. Whatever the molecular reason this can be considered a “potentiation” of the effect of RA on PA because it provokes the exponential amplification of mRNA abundance [8]. Finally, our study indicates that stabilities in mRNAs and, especially proteins, seem to play minor roles in gene regulation in general (as previously stated by [9,60]). Nevertheless, they may play important roles for some genes [12] or during dynamic processes, such as cell differentiation (discussed in [4,72]).

As the four studied organisms showed 6VP gene clusters that are statistically-enriched in genes by behaving as specific functional categories, we reaffirm the conclusion that we drew in our previous study on *S. cerevisiae*: all kind of cells use common expression strategies for the genes acting in the same physiological pathways. However, CES are not always identical between organisms, which suggests that evolution has adapted the particular expression strategies to the life styles and cell organizations of different species.

These differences in CES between species prompted us to employ them a quantitative parameter to classify organisms and to make 6VP phenograms. We are aware that a phenogram based on four organisms provides limited information because it has fewer possible topologies. In the future it would be necessary to use more organisms to draw more in-depth conclusions. In any case, the topology of the 6VP phenogram is identical to the phylogenetic tree obtained by the sequence comparison of 16-18S rRNA genes or from whole genome content [80,81]. This demonstrates the phylogenetic consistency of our functional clustering, although the relative distance between the prokaryote and eukaryotes (compared to the internal distances between eukaryotes) is clearly shorter than in sequence-based phylogenetic trees. This can reflect that DNA sequences have diverged much more than gene functions and gene regulatory mechanisms [82]. A previous study [83] constructed a phenetic dendogram based on antibiotic resistance, and concluded that it should be seen as the outward manifestation of the evolutionary history of the translational apparatus. In the same line, we conclude here that our 6VP dendogram is a global meter of the evolutionary history of whole cell functions. However, we recognize that the results from the bacterium *E. coli* are not as relevant as those obtained from the three eukaryotes, but we think that this is due mainly the poor quality GO annotation, which lowered the number of analyzable GO terms. In any case, we use *E. coli* as an outgroup for the comparative analyses of eukaryotes.

When comparing the three eukaryotes, namely two free-living yeasts separated by about 330-600 My of evolution [18,20,21] and a multicellular higher eukaryote separated from yeasts by about 1000-1100 My [18,20], the 6VP dendograms illustrate the respective influence of gene sequence evolution (nucleotide or amino acid conservation) and gene expression evolution (similarity of 6VP). Once again, the distance between HeLa and yeasts is relatively shorter than that predicted upon the evolutionary distance basis. In fact the high success rate in the systematic functional replacement of essential yeast genes with their human counterparts [82] has already suggested that gene functions and regulatory mechanisms are more conserved than gene sequences. As for human cells, it is clear that HeLa cells represent a special kind of cells, which useful for performing omics studies, but they are not representative of all kinds of human cells, especially for those with a much lower, or even zero, growth rate. Our study, thus, reflects, the similarities between eukaryotic cells growing at their fastest capability.

Regarding the similarity of the three eukaryotes in different cell components and major physiological functions, it is interesting to see how our 6VP phenogram analysis (Fig. 5B) suggests that some components and functions have closer gene expression strategies than others. As we show the identical scaling for both UPGMA and NJ trees, it becomes evident how some broad GO groups show globally less 6VP differences in the three organisms than others. Thus in the major macromolecular synthesis related to the Central Dogma, it would seem that translation has more regulatory strategy similarity than replication or transcription. It is necessary to point out, however, that GO categories have been established arbitrarily by human curators, and it not easy to be sure about the homogeneity of them all being identical. For instance, our transcription broad group includes many GO terms related to the transcription process, including transcription factors, RNA polymerases, chromatin modifiers and others, which are perhaps broader than the GO terms included in our “cytoplasmic translation” group. In any case, we think that this kind of analysis may open a new window to investigate the evolution of cellular functions.

The two yeasts are the closest related organisms based on 6VP similarity (Figure 5). It has been argued that they are almost as different from one another as from animals [18]. Yet in spite of this statement, the evolutionary time distance between them is about half that which both have as regards animals [18,20]. Moreover, as they have very similar lifestyles [19], a convergence in gene regulatory strategies would seem logical. In spite of being separated from the common ancestor with humans at the same time, globally gene expression strategies are closer to HeLa in *S. cerevisiae* than in *S. pombe*. This was unexpected because has often been stated that *S. pombe* is more similar to higher eukaryotes as the cell cycle is more similar, has many more introns than budding yeast and an RNAi mechanism, although it has no well-developed peroxisomes and proliferates mainly in the haploid state, whereas *S. cerevisiae* has peroxisomes and, in the wild, it proliferates as diploid [21]. However, it is necessary to point that our study deals with a different matter, gene regulatory mechanisms, which could have evolved to converge between budding yeast and human cells. This suggestion can be supported by the hypothesis that *S. pombe* is a more “ancient” yeast than *S. cerevisiae* based on its biological features because it appears to have undergone fewer evolutionary changes since diverging from their common ancestor [21].

We conclude that the comparative analysis of all the variables affecting the flow of gene expression is a useful strategy for investigating the regulatory strategies used by living cells, at least by eukaryotes. We also conclude with our study that the transcription rate is the main determinant of the amount of the corresponding protein, and that all kinds of organisms, either prokaryotic, single-cell or higher eukaryotes cells, use CES for the genes acting on the same physiological pathways. This feature can be reflected as a 6VP that defines the average behavior of a given gene group. CES are more clearly seen for the genes coding for large and stable protein complexes, such as the ribosome or the spliceosome, but can be seen even in groups of genes that do not form stable complexes, but are functionally related, like those involved in different metabolic pathways. Some of the 6VP are similar between different organisms, which reflects either a common evolutionary origin or a convergent evolution due to similar functional constraints or horizontal gene transfers. We propose that comparing 6VPs for a series of organisms is another way to draw phenograms to reveal how the environment of cells influences gene expression strategies. The use of omics data for phenetic classifications is not new [74,83,84], but our analysis extends the available tools for phenetic classifications by allowing to study global expression strategies adapted to lifestyle.

## Abbreviations

GO: gene ontology
TR: transcription rate
RA: mRNA amount
RS: mRNA stability
PA: protein amount
TLRi: individual translation rate
PS: protein stability

## Data Availability

All the data generated or analyzed during this study are included in this published article [and its Supplementary Information files].

## Acknowledgments

The authors are especially grateful to Julen Mendieta, who participated in the initial analyses of this work, and to all the GFL laboratory members for discussion and support, to G. Ayala for his help in the initial statistical analyses and to Drs. J. Peretó and F. González-Candelas for critically reviewing the manuscript.

## Authors’ contributions

JEP-O. and JG-M conceived the study, and analyzed and interpreted the data. JF-M, A.F-D and JG-M performed the bioinformatics and statistical analyses. JEP-O wrote the paper. All the authors revised and approved the paper.

## 9. Supplementary Figure legends

**Figure S1.-Additional clusters from the four studied organisms showing defined profiles and statistically enriched GO terms.** See Figure 3B for details.

## 10. Supplementary Tables

**Table S1. List of the genomic data for the six variables: RA, TR, RS, PA, TLRi, PS used for Pearson’s and Spearman’s correlations in the four organisms.**

**Table S2. List of the genomic data for the six variables in rank order, RA, TR, RS, PA, TLRi, PS, used for 6VP construction, clustering and phylogenetic trees in the four organisms**.

**Table S3. List of macromolecular complexes and GOs not belonging to the macromolecular complexes used for the Figure 4 comparison**.

**Table S4. List of the GO terms used for the phenogram reconstruction in Fig. 5**

## 11. Appendices

**Appendix 1. Whole set of clusters obtained for *S. cerevisiae*.**

**Appendix 2. Whole set of clusters obtained for *S. pombe*.**

**Appendix 3. Whole set of clusters obtained for *E. coli*.**

**Appendix 4. Whole set of clusters obtained for HeLa.**

